# Massive perturbation of sound representations by anesthesia in the auditory brainstem

**DOI:** 10.1101/2024.02.06.579173

**Authors:** Etienne Gosselin, Sophie Bagur, Brice Bathellier

## Abstract

Anesthesia modifies sensory representations in the thalamo-cortical circuit, but is considered to have a milder impact on peripheral sensory processing. Here, tracking the same neurons across wakefulness and isoflurane anesthesia, we show that the amplitude and sign of single neuron responses to sounds are massively modified by anesthesia in the cochlear nucleus of the brainstem, the first relay of the auditory system. The reorganization of activity is so profound that decoding of sound representation in anesthesia is not possible based on awake activity. However, population level parameters, such as average tuning strength and population decoding accuracy are weakly affected by anesthesia, explaining why its effect has previously gone unnoticed when comparing independently sampled neurons. Together, our results indicate that the functional organization of the auditory brainstem largely depends on the network state and is ill-defined under anesthesia. This demonstrates a remarkable sensitivity of an early sensory stage to anesthesia, which is bound to disrupt downstream processing.

**Teaser:** Anesthesia compromises the normal transmission of sensory information as early as the first relay in the auditory system.

## Introduction

Modern sensory neurophysiology has shifted towards the use of awake animals to avoid the effects of anesthetics on neural response properties (*1*–*5*) which appear much larger than previously thought in the thalamic and cortical stages (*6*, *7*). However, this shift of experimental practice is mostly focused on processing higher stages including cortex (*8*–*10*), thalamus (*11*–*13*) and colliculus (*14*, *15*). For early sensory areas, located deep in the brainstem, most of our knowledge still relies on recordings performed under anesthesia based on the assumption that this has a minimal impact on sensory processing at this stage (*16*–*21*). Recent progress of extracellular recording and even imaging (*22*) techniques now give access to the brainstem in awake conditions, including pioneering recordings in the cochlear nucleus, the first relay of the auditory system (*22*–*24*). These studies suggest that sound tuning properties in the awake early auditory system are similar to those observed under anesthesia despite a clear reduction of spontaneous activity and a small increase of response threshold under anesthesia, in line with early observations in decerebrate and paralyzed cats (*25*, *26*). However, these conclusions are not based on the direct comparison of sound representations in the same neurons across states, which is necessary to determine if the sensory code is preserved one-to-one under anesthesia (*6*). Here, we performed extracellular recordings with a linear multielectrode array (Neuropixels 1.0) (*27*) in the cochlear nucleus of mice across wakefulness and isoflurane anesthesia. We observed a drastic modification of spike statistics under anesthesia, with a strong reduction of spontaneous firing rates. This heavily impacted the performance of template-based spike sorting algorithms. Tailoring the algorithms to circumvent this issue revealed that anesthesia modifies the functional identity of single neurons well beyond the effects previously described. We observed that many neurons that were weakly or non-responsive in the awake state become strongly sound-driven in anesthesia, and vice-versa. These bidirectional changes are so profound that population decoders of sound identity trained on responses observed in one state could not decode sound identity in the other state. In line with this, we observed that sound representations occupy different subspaces of neural activity in wakefulness and anesthesia. However despite these massive changes, sound representations are similarly accurate in both states. This explains why previous observations in distinct datasets failed to identify the profound changes that became apparent when we tracked neuronal identity. By changing the neurons carrying the representation of sounds, isoflurane unavoidably modifies the integration of information downstream in the auditory system which likely contributes to the reduction of sound-related information observed in auditory cortex responses under anesthesia(*6*).

## Results

### Tracking neurons across wakefulness and anesthesia requires fine tuning of spike sorting methods

To access the cochlear nucleus in awake mice, we trained mice to stay quietly head-fixed held by a head-post previously positioned for reliable stereotaxic placement in the electrophysiology apparatus. Thanks to this preparation, Neuropixels 1.0 probes could be inserted at a predefined angle and entry point (**Fig. 1A, B,** see Methods). Localisation of the probe was fine-tuned through repeated penetrations until time-locked responses to sounds could be detected. Recording could be repeated up to 3 days in a row, allowing us to obtain 7 recording sessions in 4 mice. After the last session, the exact positioning of the probe was determined with post-hoc histological identification of the electrode tracks marked with fluorescent dyes (**Fig. 1B**). 5 recordings targeted the postero-ventral cochlear nucleus (PVCN) and 2 the dorsal cochlear nucleus (DCN). Activity in the cochlear nucleus was recorded during presentation of a broad set of sounds (307 short sounds, duration <500ms, Pure tones, amplitude or frequency modulations and complex sounds, **Fig. 1C**, and ten 30s natural sounds), first in the awake state (80 min session), then under 1.1% isoflurane anesthesia corresponding to a light narcosis state (**Fig. 1D, E**). Strikingly, visual inspection of raw extracellular electrophysiology traces was sufficient to observe a clear switch of brain activity regime between wakefulness and anesthesia based on the marked amplitude drop of rapid fluctuations (**Fig. 1F, G**). This could be quantified by computing the mean deviation of high frequency fluctuations (high-pass filtering above 300 Hz) which decreased two- to four-fold (3.6 ± 0.4) under anesthesia outside of sound presentations (**Fig. 1H**). The magnitude of high frequency fluctuations, sometimes quantified as multiunit activity, corresponds for the most part to the summation of extracellular action potentials and has been found to scale sublinearly with the local population firing rate (*28*). Therefore, we can estimate that the population firing rate decreases more than 3-fold between wakefulness and anesthesia, in line with previous reports (*25*, *29*–*31*).

**Figure 1:**
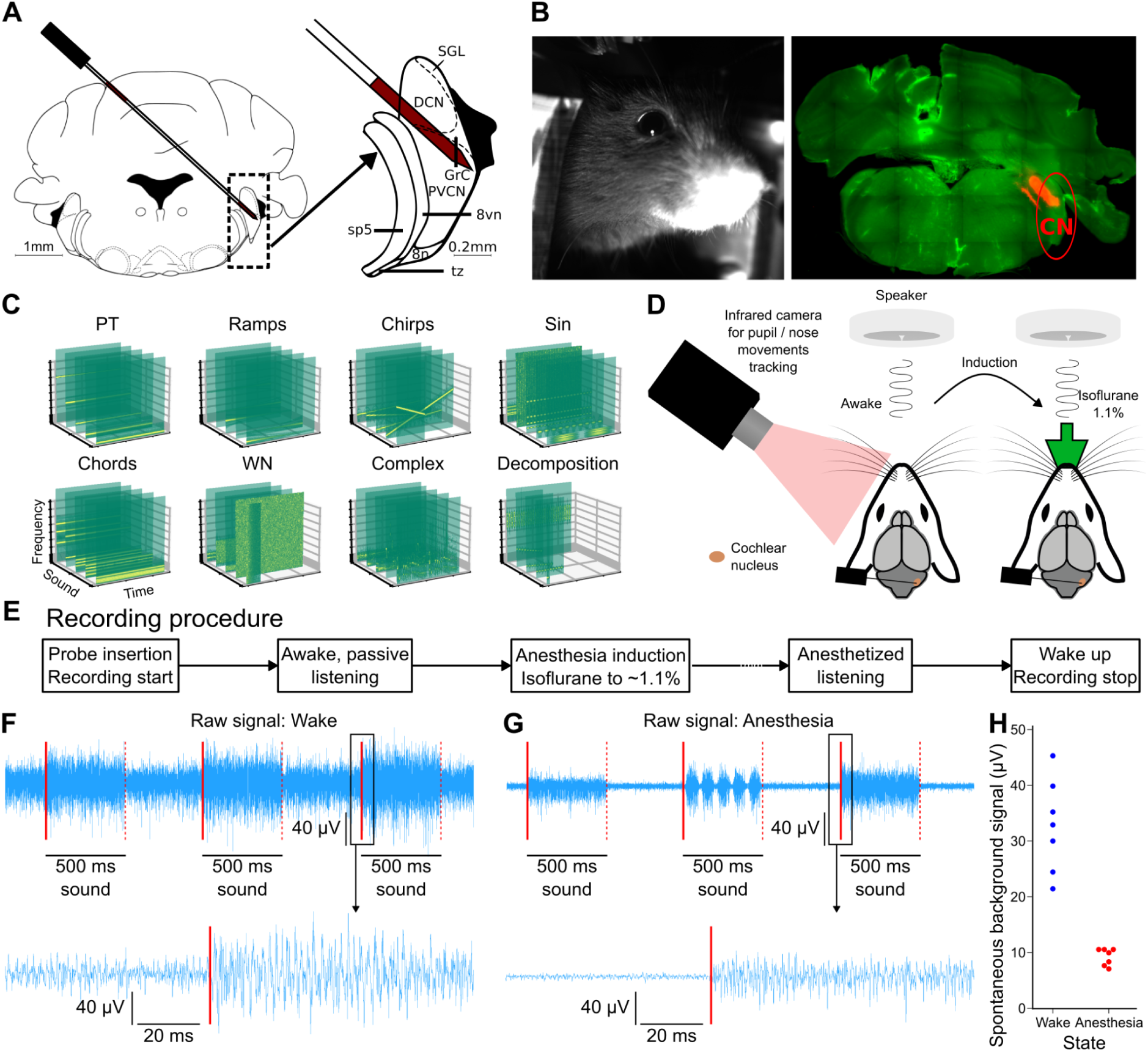
Electrophysiology recordings of cochlear nucleus neurons in the awake and anesthetized mouse. **A.** Schematic of probe insertion with a zoom on the cochlear nucleus area (right). **B.** Mouse state was monitored with an infrared camera showing the face of the mouse (left). Targeting is verified with histology showing the fluorescent probe tract (right). **c.** Spectrogram representation of 5 example sounds from each of the 8 categories included in the sound set. From left to right, top to bottom: pure tones, linearly amplitude ramping sounds, linear frequency modulations, sinusoidal amplitude modulations, sums of pure tones, broadband and filtered noises, complex natural sounds and component decomposition of complex sounds. **D.** Schematic drawing of the experimental setup. **E.** Sketch of the experimental procedure. A speaker placed in front of the head-fixed, awake mouse delivers the sounds for a duration of 80 minutes. The mouse is then anesthetized and the sounds are delivered again. **F.** 3 seconds snippet of signal from 1 contact after bandpass filtering and common-average reference preprocessing (top), and 150 millisecond zoom (bottom) during the awake procedure. **G.** Same as **F** during the anesthetized procedure. **H.** The magnitude of high frequency fluctuation (median average deviation estimate of the standard deviation of the signal) during spontaneous activity is lower under anesthesia (p= 0.01, n= 7 recording session, Wilcoxon ranksum test).

This massive change of activity regime had a strong impact on spike sorting performance to extract single unit activity. Applying the spike sorter Kilosort 2.5 on the full concatenated wakefulness and anesthesia datasets, we observed that much fewer spikes were detected under anesthesia as expected from the drastic decrease in spiking activity (**Fig. 2A, B**). However, visual inspection of the data revealed that many clearly visible spike waveforms were not assigned to any clusters during anesthesia (**Fig. 2B**). Similar results were obtained with other spike sorters tested though the SpikeInterface software (*32*). This poor performance of out-of-the box spike sorting methods is likely attributable to the large change in spike statistics, including the temporal proximity of spikes and the level of concurrent activity, two factors known to significantly impact spike sorting (*33*–*37*). We therefore tailored two complementary methods to optimize tracking neurons between states (**Fig 2C**). Our first approach consisted of running spike sorting separately on each state, independently optimizing the quality of units and then matching the detected units between states based on comparison of their waveform similarity to a chance distribution (see Methods). In total, this Split spike sorting (Split s.s.) led to 173 (anesthesia) and 163 (wake) single units respectively. This method was able to detect anesthesia spikes missed when sorting on the full dataset (**Fig. 2B**). It also resulted in more similar numbers of isolated spikes (**Fig. 2A**) and units between wakefulness and anesthesia because the lower background noise under anesthesia likely compensated for the increased activity in wakefulness. Strikingly, only 18 of these were matched between the two states (**Fig 2D, E**, see waveforms of matched neurons in **Supplementary Fig. 1**). This suggested that different neurons are active in the awake or anesthetized state but showed that a subset was sufficiently active in both states to be tracked with confidence. The stringent criteria of waveform matching likely underestimates the true number of trackable neurons since some drift is expected over the two hour recordings (*38*). Our second approach aimed at maximizing the number of neurons directly tracked between states. A key step in the Kilosort 2.5 algorithm is the initial estimation of waveform templates which are then used to associate individual spike waveforms to clusters. Given the drop in firing rate during anesthesia, this estimation step was dominated by awake spikes and the templates were likely biased towards waveforms typical of the awake state. We therefore forced the algorithm to define initial templates on spikes from anesthesia data, during which the low level of background activity facilitates clean template estimation. We then applied looser parameters during template matching of the full session to allow the algorithm to match spikes despite shifts in waveform amplitude. This Consensus spike sorting (Consensus s.s.) tracked 163 single units between states with high waveform similarity across states (**Fig. 2E**, see sample waveforms across states in **Supplementary Fig. 2**). In wakefulness, fewer spikes were successfully clustered (**Fig. 2A**), validating the idea that certain neurons fall silent during anesthesia. Moreover, the clusters were noisier than those identified in the stringent split spike sorting. This could be directly measured by comparing the reliability of their response to multiple presentations of the same sound. Clearly, response reliability was higher in units extracted by the split spike sorting than the consensus spike sorting (**Fig. 2F**). In conclusion, despite the difficulty of spike sorting across wakefulness and anesthesia, that may explain the lack of studies directly comparing the two states, we robustly tracked the activity of the same single units. This yielded two datasets, one with high reliability but less units tracked across states and the other with a larger number of less reliable units.

**Figure 2:**
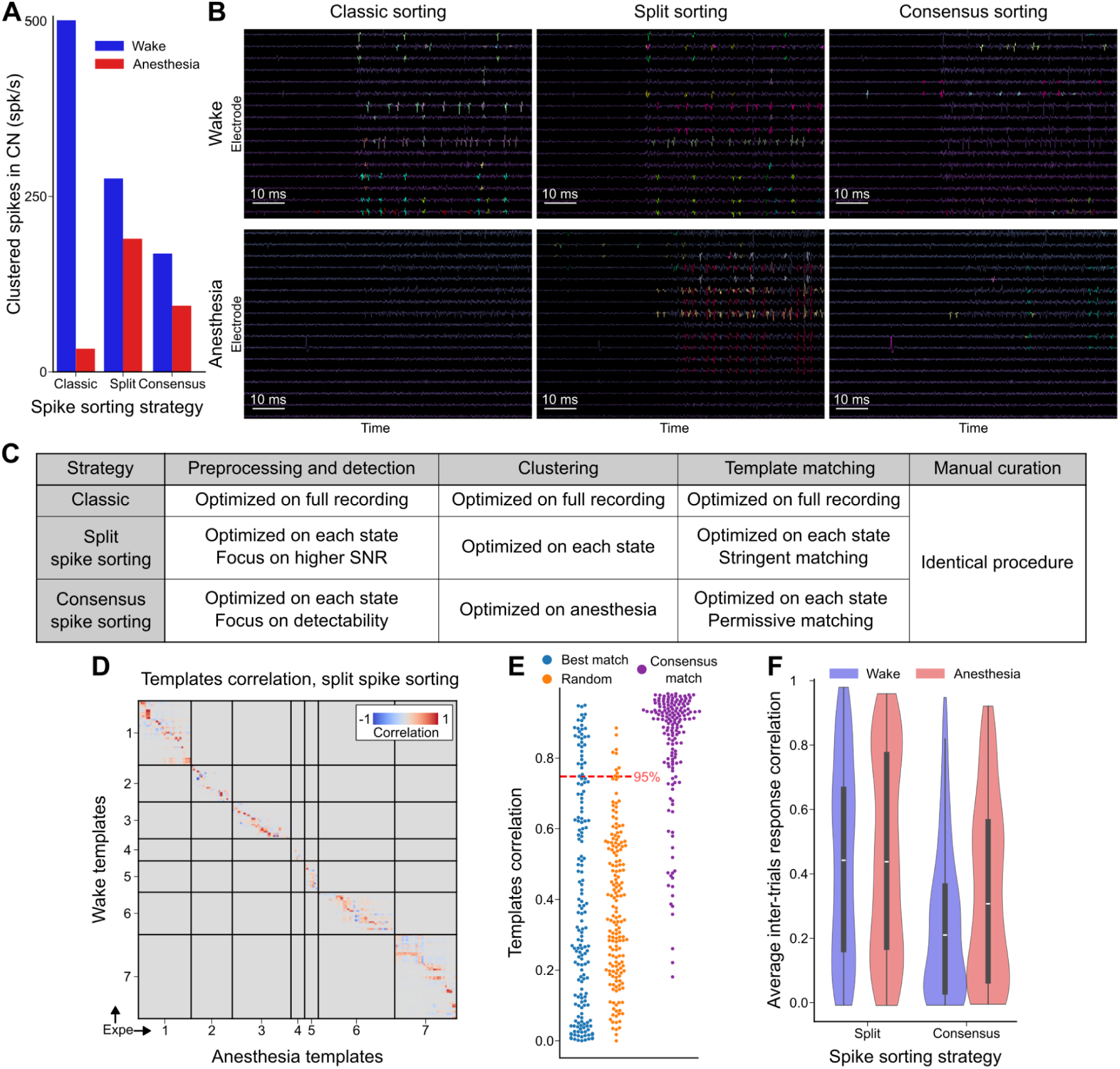
The broad activity changes induced by anesthesia impact spike sorting. **A.** Number of spikes per second assigned to single-unit clusters for classic, split, and consensus spike sorting strategies, in each state. **B.** Phy GUI visualization of clustered spikes in both states for the three spike sorting strategies. Each color represents spikes assigned to one cluster. **C.** Comparative table of the 3 spike sorting strategies for the main steps of Kilosort 2.5 and Phy GUI spike sorting procedure. **D.** Correlation matrix between spike templates of neurons defined during wake and anesthesia (split s.s.). Correlations between templates of neurons from different recordings are manually set to 0. **E.** Value of correlation with the best correlated anesthesia neuron template for each wakefulness neuron template, restricted to neurons from the same recording (in blue, split s.s.). Value of correlation with the best correlated anesthesia neuron template for each wakefulness neuron template, restricted to neurons from different recordings, equivalent to a random distribution (in orange, split s.s.). Value of correlation between the template defined on wakefulness and the template defined on anesthesia for all neurons (in purple, consensus s.s.) displayed as comparison. **F.** Average correlation of temporal response of neurons between multiple presentations of each sound for all neurons in both datasets. This provides a measure of response reliability which is significantly lower for consensus spike sorting (Wake: p=6e-10, N=163, anesthesia: p=1e-3, N=163 for anesthesia, Mann-Whitney U test).

### Anesthesia strongly decreases spontaneous activity and weakly increases intensity thresholds and tuning precision

To validate these two spike-sorted datasets, we verified that the previously observed effects of isoflurane anesthesia on the cochlear nucleus are present in our data, when treating awake and anesthetized states as two separate datasets. Several studies have compared, at the population average level, a number of functional properties across datasets recorded separately under anesthesia and in wakefulness. They found a clear drop of spontaneous firing rates under anesthesia which led to a disappearance of the negative responses to sounds that are clearly visible in wakefulness (*25*, *30*). These studies also reported narrower tuning width and an elevated intensity threshold for pure tone responses (*25*, *30*). The drastic decrease of spontaneous activity when no sound was presented was easily observable in our raster plots (**Fig. 3A-D**). To quantify it, we averaged the spontaneous firing rate of each single unit across all the ∼2s intervals which separated blocks of sound stimulation (see **Fig. 3B, D**). This amounted to ∼4 minutes of spontaneous activity. Spontaneous firing rates ranged between 0 spikes per second (spk/s) and ∼100 spk/s, with a shift of the distribution towards low firing rates under anesthesia. 69% (Split s.s.) and 51% (Consensus s.s.) of the units had a spontaneous firing rate above 1 Hz in the awake state, and only 20% (Split s.s) and 16% (Consensus s.s.) under anesthesia (**Fig. 3E**). In comparison, only 1% (Split s.s.) and 2% (Consensus s.s.) of the units had zero spontaneous spikes in wake, while 47% (Split s.s.) and 32% (Consensus s.s.) of the units were completely silent in anesthesia. These observations are fully consistent with previous reports and with the drastic decrease of spontaneous neural firing (**Fig. 1F, H**). Contrariwise, in the non-auditory brainstem regions neighboring the cochlear nucleus, spontaneous activity levels increased in anesthesia indicating that isoflurane produces differential dynamical effects on different brain structures (**Fig. 3A, C**). Reduced spontaneous activity under anesthesia led to the disappearance of sound-driven firing rate reductions (**Fig. 3A, C**). 20% (Split s.s.) and 15% (Consensus s.s.) of the single units displayed an average firing rate decrease of more than 1 Hz in wakefulness and only 1 % (Split s.s.) and 3% (Consensus s.s.) under anesthesia (**Fig. 3F**). Despite the massive change in the spontaneous network dynamics, positive responses to sounds were still clearly visible under anesthesia (**Fig. 3A-D**). To characterize these responses, we measured the preferred frequencies of each neuron and its tuning width for pure tones at 50 and 70 dB SPL. Anesthesia slightly reduced the occurrence of tuning to higher frequencies although not significantly (**Fig. 4A**, p=0.08 for split s.s., p=0.27 for consensus s.s., Kolmogorov-Smirnov test for distinct distributions). The half-width of tuning curves at 50 and 70dB SPL (**Fig. 4B**), revealed, as previously observed (*30*), that tuning is slightly broader in the awake state than under anesthesia (**Fig. 4C**). This effect was larger at lower intensity suggesting that the improved tuning width may be related to an increased sound level threshold. To quantify this, we measured response delays for pure tones ramped in intensity from 50 to 70 dB over 500 ms (**Fig. 4D**). Units responded on average at a lower sound level (i.e. earlier in the ramp) in the awake than in the anesthetized states (**Fig. 4E**), consistent with an increased intensity threshold, which we could observe here only for the high-intensity range of thresholds (>50dB).

**Figure 3.**
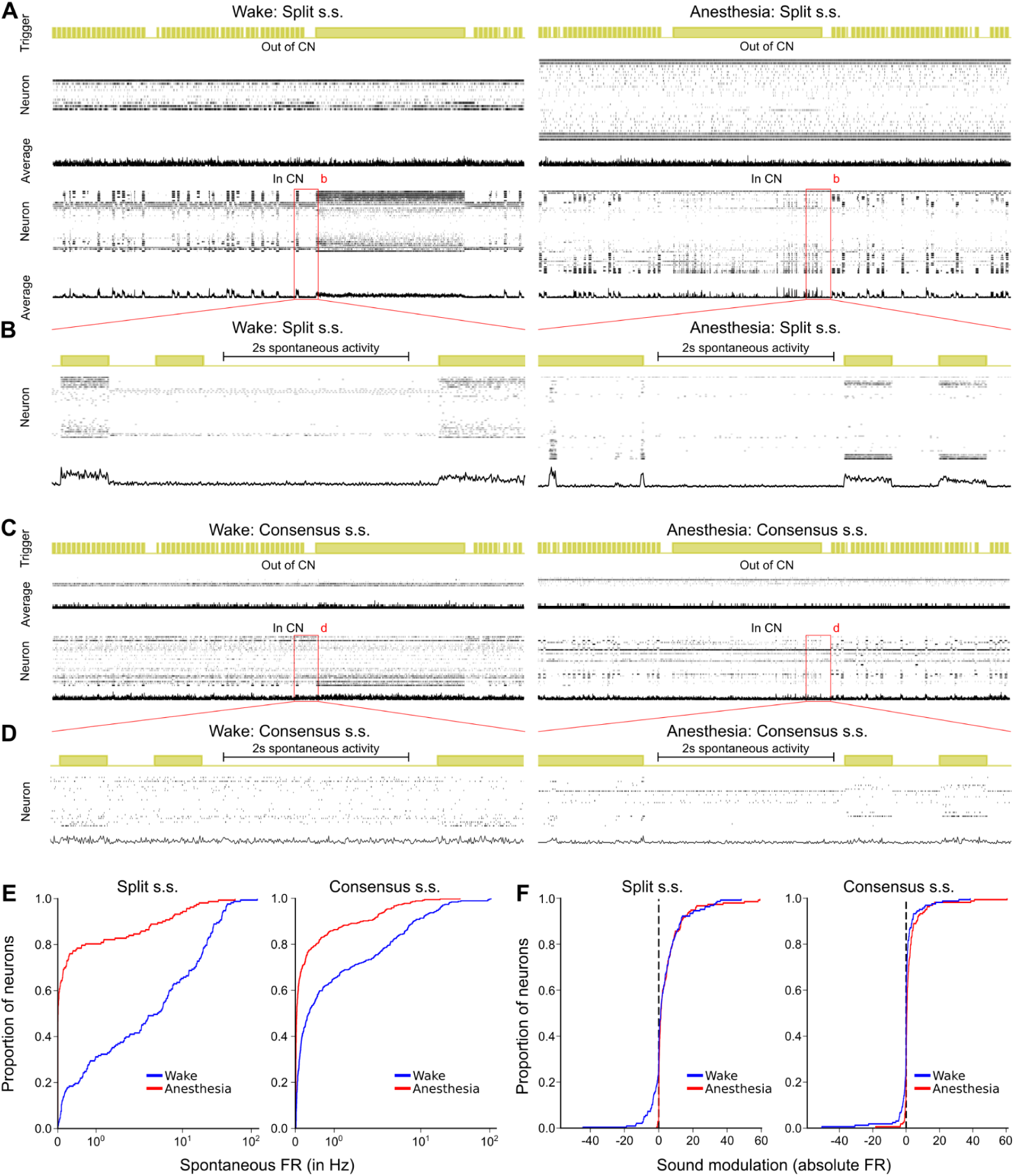
: Anesthesia reduces spontaneous firing and inhibitory responses to sounds. **A,C.** Example raster plots for 100s of data during awake (left) and anesthetized (right) procedures, for split s.s. (**A**) and consensus s.s. (**C**). Sound presentations are displayed in yellow (Trigger). Rasters of neurons (Neuron) and the population-average of spikes (Average) are plotted for neurons included in the dataset based on depth boundaries (In CN) and for left-out neurons (Out of CN). **B,D.** Same as **a,c**. for the duration defined by the respective red areas during awake (left) and anesthetized (right) procedures. **E.** Cumulative distribution function of the absolute spontaneous firing rate of neurons (concatenated data extracted from the spontaneous activity recorded between blocks of presentation of sounds) for wakefulness (blue) and anesthesia (red) and for both split and consensus spike sorting (Split s.s.: p=1e-29, N=163. Consensus s.s.: p=4e-31, N=163, Kruskal-Wallis test). **F.** Cumulative distribution function of the average modulation of baseline subtracted firing rate of neurons in response to sounds during both procedures (Split s.s.: p=0.04, N=163. Consensus s.s.: p=3e-6, N=163, Kruskal-Wallis test).

**Figure 4:**
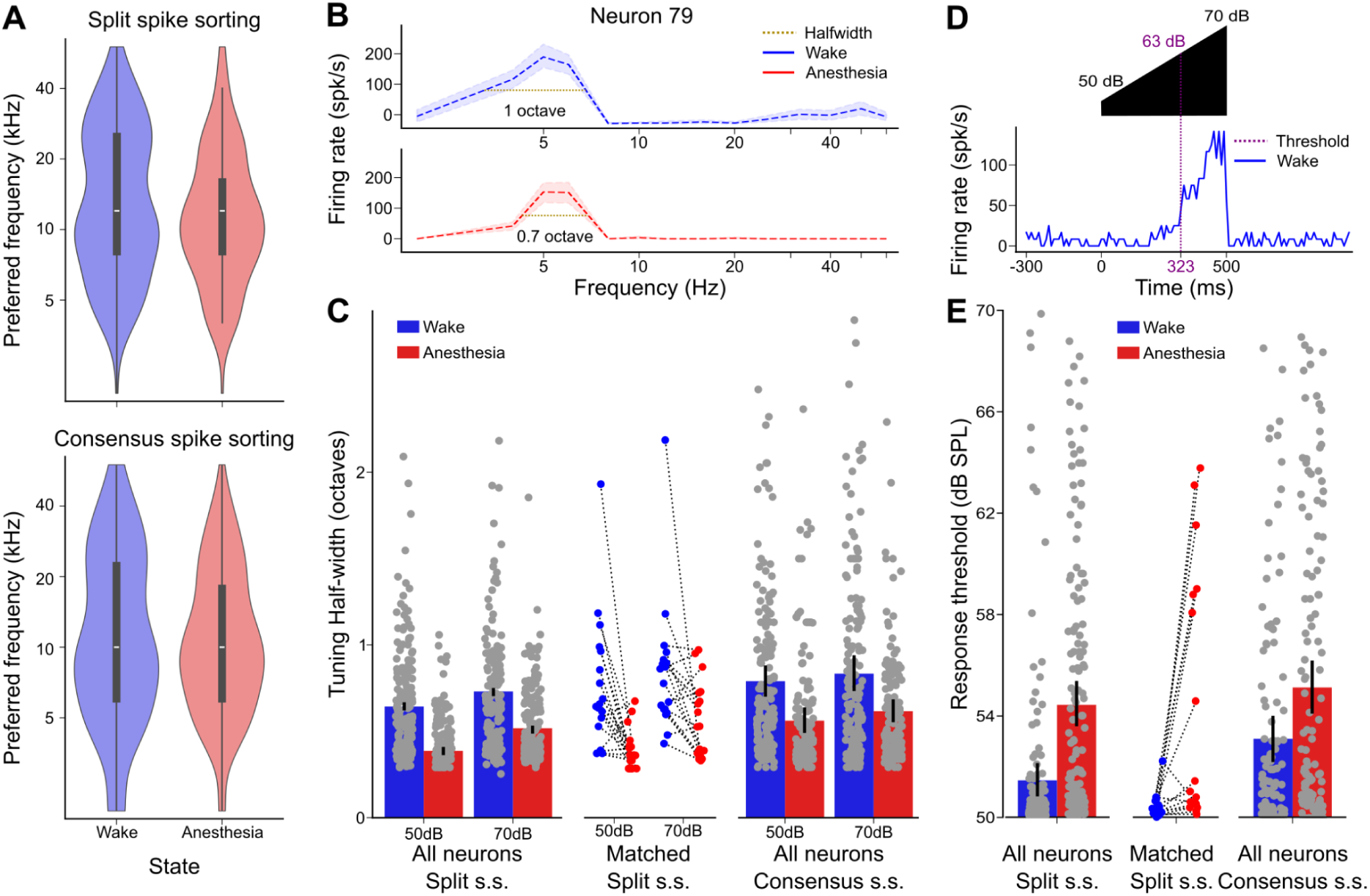
Anesthesia sharpens frequency tuning and increases intensity threshold. **A.** Distribution of the best frequencies of neurons for 70dB pure tones for both procedures, computed as the maximal average response. **B.** Example tuning curve for one neuron displaying the half-width in each state. **C.** Barplots of the tuning halfwidths (in octaves) for both procedures (mean±SEM, Mann-Whitney U test for difference in distributions: Split s.s. 50dB = 0.58±0.05 for N=172 wake neurons, 0.51±0.05 for N=132 anesthesia neurons, p = 1e-5. Split s.s. 70dB = 0.82±0.05 for N=173 wake neurons, 0.52±0.05 for N=162 anesthesia neurons, p =1e-8. Matched split s.s. 50dB = 0.81±0.09 for N=17 wake neurons, 0.52±0.08 for N=14 anesthesia neurons, p =5e-3. Matched split s.s. 70dB = 1.1±0.15 for N=18 wake neurons, 0.57±0.05 for N=18 anesthesia neurons, p =1e-3. Consensus s.s. 50dB = 0.80±0.04 for N=158 wake neurons, 0.64±0.04 for N=142 anesthesia neurons, p =4e-4. Consensus s.s. 70dB = 0.93±0.06 for N=159 wake neurons, 0.73±0.04 for N=162 anesthesia neurons, p =1e-2). **D.** Example response of one neuron to an up-ramping sound at its best frequency, displaying the response start value. The threshold in amplitude is calculated as the intensity of the sound at the time when the response exceeds twice the standard deviation of baseline activity, at the best frequency of the neuron. **E.** Barplots of the amplitude threshold of response for all neurons during presentation of a pure frequency increasing linearly in intensity (mean±SEM, Mann-Whitney U test for difference in distributions: Split s.s. = 52.4±0.4 for N=114 wake neurons, 55.1±0.5 for N=135 anesthesia neurons, p = 6e-9. Matched split s.s. = 51.8±1.2 for N=13 wake neurons, 53.7±1.0 for N=16 anesthesia neurons, p = 4e-2. Consensus split s.s. = 53.1±0.4 for N=118 wake neurons, 55.1±0.5 for N=130 anesthesia neurons, p = 2e-3).

### Isoflurane anesthesia massively modifies which neurons contribute to the sound representation

Intrigued by the contrast between the strong impact of anesthesia on spontaneous activity (**Fig. 3**) and its weak impact on responses to sounds (**Fig. 4**), we further explored sound representations across states by plotting single cell tuning profiles for units that could be tracked both in the anesthetized and awake states (**Fig. 5A, B**). For both split or consensus spike sorting, we observed that the changes of response properties at the single cell level were more drastic than suggested by average population properties. First, we observed units that were only responsive in one of the two states: some units with salient pure tone tuning curves in the awake state displayed hardly any response under anesthesia (e.g. **Fig. 5A**: Neurons 2 & 3; **Fig. 5B**: Neurons 6), and conversely certain unresponsive units in the awake state developed robust pure tone tuning under anesthesia (e.g. **Fig. 5A**: Neurons 11 & 14; **Fig. 5B**: Neurons 135 & 3). We also identified changes in the sign of the response with units inhibited by sounds in the awake state and excited under anesthesia. This held both for cells that were broadly inhibited across sound frequencies (e.g. **Fig. 5B**: Neurons 1), and those with narrowly tuned inhibition (e.g. **Fig. 5A**: Neuron 7). Consistent with the low spontaneous activity in anesthesia, only a very low number of neurons developed inhibitory responses in this state (e.g. **Fig. 5B**: Neuron 28). Overall, the response levels for a given sound clearly changed across the two states, but some neurons had weaker variations (e.g. **Fig. 5A**: Neurons 10, 12 & 13; **Fig. 5B**: Neurons 14, 73 & 79). When a preferred frequency could be identified both in wakefulness and anesthesia, it was usually similar across states (**Fig. 5A, B**). Hence, anesthesia primarily impacts whether or not a neuron participates in the representation but not which frequency it represents. Systematic quantification of response magnitudes at best frequency for all units identified in both states (**Fig. 5C, D**) confirmed large bidirectional cross-state variations observed in sample neurons **Fig. 5A, B**. Strikingly, because variations were bidirectional, the overall distribution of responses amplitudes was similar in both states (**Fig. 5C, D**), except for the lack of negative responses under anesthesia, consistent with **Fig. 3F**. Together these observations indicate that despite a similar frequency coverage and a similar distribution of response in wakefulness and anesthesia, isoflurane had a massive effect on which neurons participate in the sound representation. As a result, a given sound does not activate the same pattern of cochlear nucleus neurons in wakefulness and under anesthesia.

**Figure 5:**
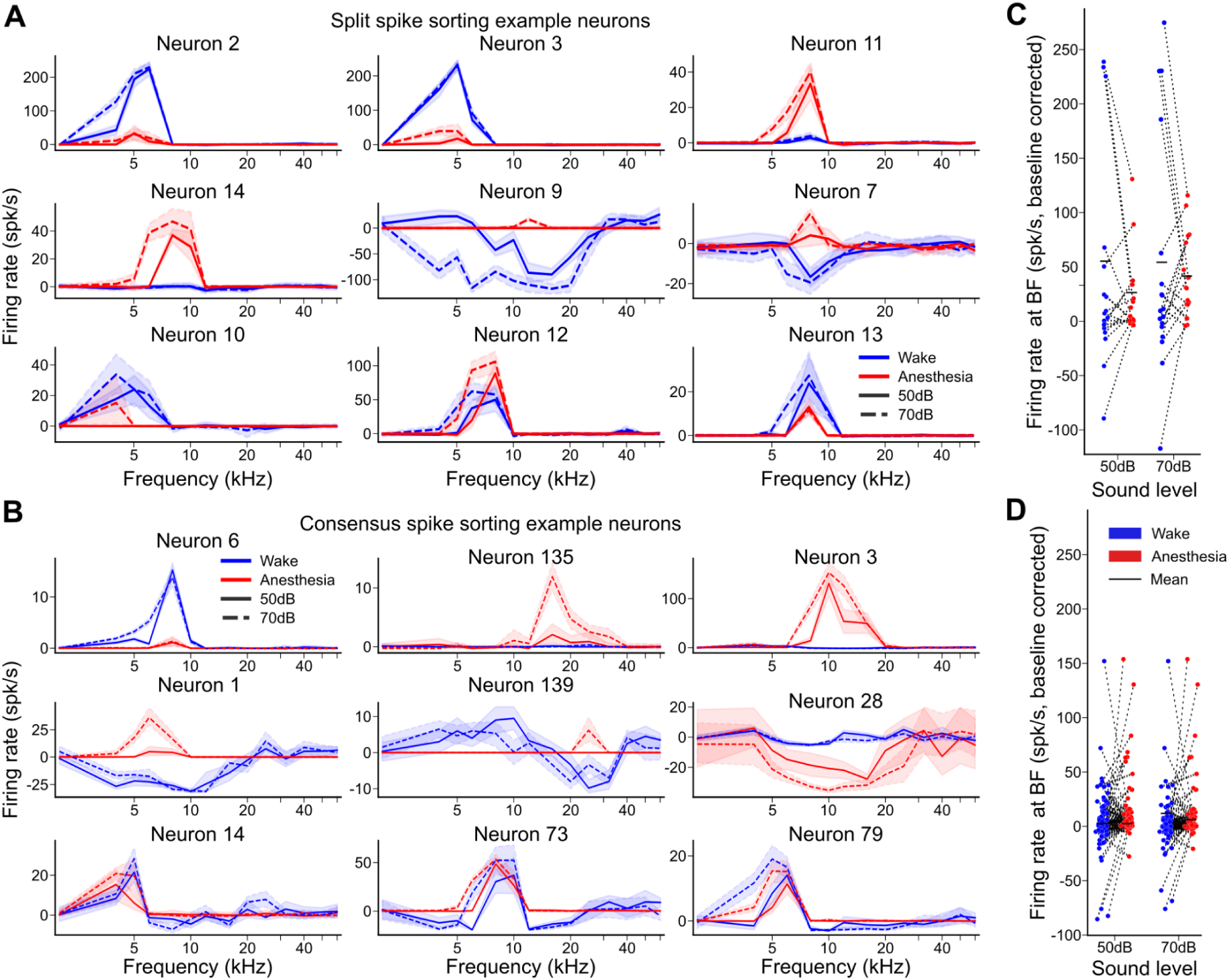
Massive modulation of sound responses in neurons tracked across states. **A.** Pure tone tuning curves for 9 single units whose spike waveform could be tracked across states (awake = blue and anesthetized = red) for split spike sorting. The dashed and plain lines represent the tuning at 50 and 70 dB SPL respectively. **B.** Same as a. but for consensus spike sorting. **C.** Baseline-corrected firing rate for neurons at their best frequency during both states for split spike sorting. **D.** Same as c. but for consensus spike sorting.

### Anesthesia shifts the sound representation in a different neural population subspace

We reasoned that this reorganization of sound-driven activity must have consequences for the downstream decoding of cochlear nucleus activity. To quantify these effects and estimate the amount of sound information carried by the cochlear nucleus across states, we used cross-validated population activity classifiers trained to discriminate between sounds based on the combined activity of all recorded neurons. For generality, the classifiers used the information carried by the full activity time-course of each neuron (**Fig. 6A**) but similar results were obtained when using only firing rates averaged across the sound duration. By testing the classifier on a test set of population responses originating from the same state, we evaluated the amount of sound information in this state (**Fig. 6A-C**, same state). By testing the classifier on population responses from a different state, we quantified the decodability of neural activity across states, and thereby the impact of anesthesia on the structures downstream of the cochlear nucleus (**Fig. 6A-C**, cross state). Same-state decoding accuracy was similar between wakefulness and anesthesia for both spiking sorting strategies (0.80 ± 0.02 for wakefulness and 0.73± 0.02 for anesthesia for split spike sorting, **Fig. 6B**, 0.49 ± 0.02 and 0.60 ± 0.02 for consensus spike sorting, **Fig. 6C**, chance level: 0.0032). Note that the lower classification accuracy with the consensus spike sorting is likely due to the fraction of spikes missed with this approach. Sound classification was only slightly worse when restricting the analysis to the 18 neurons matched across states from the split spike sorting (Awake accuracy = 0.56 ± 0.02, Anesthetized accuracy = 0.49± 0.02, **Fig. 6D**), although they only represent 10% of the full dataset population. Interestingly, while information levels were similar across states, the underlying code was dramatically changed as shown by the drop of across state classification accuracy (0.08 ± 0.01 and 0.06 ± 0.01 for split, 0.10 ± 0.01 and 0.08± 0.01 for consensus spike sorting, **Fig. 6B, C**). The same effect is observed, if we restrict the classification to only the 14 pure tones at both amplitudes (**Fig. 6B, C**). This result suggests that the representations of sounds in the awake and anesthetized states populate different regions of the neural activity state space, i.e. that sounds are coded by different combinations of cochlear nucleus neurons. In order to demonstrate this, we reduced the dimension of the dataset using principal component analysis over all sound-driven activity patterns and retained for display purposes only the first three components. We then projected the population vectors representing each sound in each state into this 3-dimensional space. The resulting plots (**Fig. 6D, E)** clearly show that sound-evoked activity patterns under anesthesia and in wakefulness occupy different regions of the neural population activity space. To complement this qualitative analysis, we classified state identity using a linear support vector machine with the data projected into this 3 principal component space, whose decision boundary is displayed as a solid black line in **Fig. 6D, E**. The accuracy of decoding on a 10-fold cross-validation of the support vector classifier was above 90% for both datasets. Hence, anesthesia transfers sound representations into different subspaces of the neural population activity state-space.

**Figure 6:**
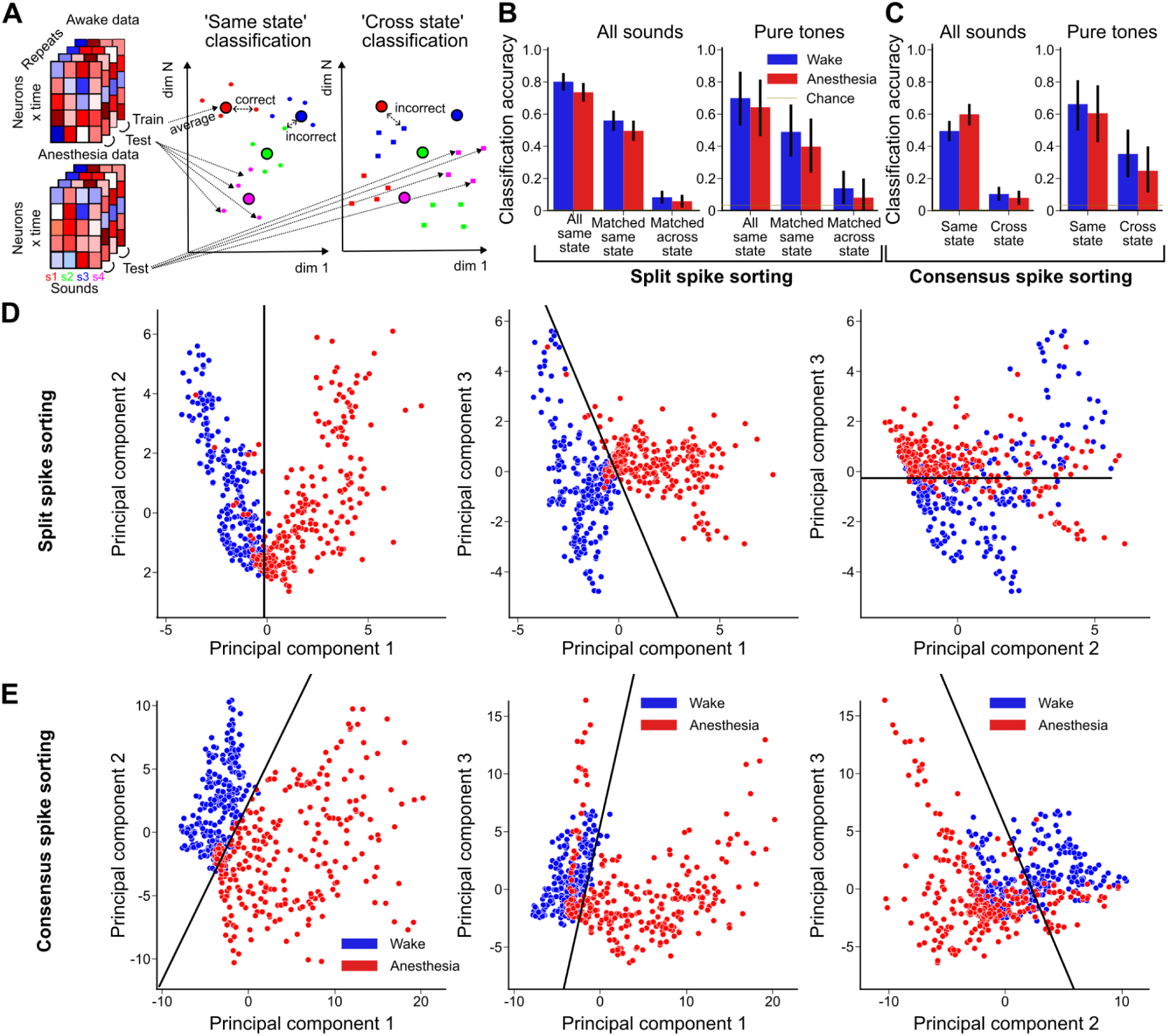
Isoflurane anesthesia redistributes sound information across the cochlear nucleus neural population. **A.** Schematic drawing of the nearest-neighbor classifier procedure for decoding within state and across states. **B.** Classification accuracy of all sounds and pure tones only by the awake and anesthetized datasets for same-state classification and cross-state classification using the split spike sorting dataset. Across state classification is much worse than within state classification (All sounds: p=5e-80, N=614. Pure tones: p=3e-7, N=28, Wilcoxon signed-rank test) **C.** Same as **B.** for the consensus spike sorting dataset (All sounds: p=8e-50, N=604. Pure tones: p=0.1, N=28, Wilcoxon signed-rank test). **D,E.** Distribution of sound responses during wake (blue) and anesthesia (red) after projection into the first three principal components for split s.s (**D**) and consensus s.s (**E**). The black line indicates the plane of best separation between the two states as defined by linear support vector machine classification.

### Isoflurane anesthesia differentially impacts the representations of distinct acoustic cues

Sound representations in the awake and anesthetized states encode similar amounts of sound information in the cochlear nucleus, however this information is carried by different codes. We therefore investigated if the structures of these two codes are similar or distinct. To answer this question, we unfolded classifier performance into confusion matrices (**Fig. 7A, B**) and quantified sound classifiers accuracy for specific categories of sounds (**Fig. 7C**. We used classifiers either on the time-averaged firing rate of each neuron (rate code) or on the full time course of their activity (temporal code). For overall classification, we observed that the temporal code led to larger decoding accuracies in both states (**Fig. 7B**; Split spike sorting: 0.74±0.02 for the temporal code, 0.53±0.02 for the rate code, p-value=2.10^-16^ for wake, 0.69±0.02 for the temporal code, 0.47±0.02 for the rate code, p-value=5.10^-17^ for anesthesia. Consensus spike sorting: 0.49±0.02 for the temporal code, 0.53±0.02 for the rate code, p-value = 0.2 in wake, 0.60±0.02 for the temporal code, 0.48±0.02 for the rate code, p-value=2.10^-5^ in anesthesia). This is consistent with previous observations that temporal features in neural activity are crucial to describe time-varying sounds in the subcortical auditory system(*39*, *40*). When quantifying accuracy for specific sound categories we observed that pure tone identification was equally efficient in wakefulness and under anesthesia for the higher sound level (70 dB SPL) but was impaired under anesthesia at lower level (50 dB SPL, **Fig. 7C**). This likely reflects the higher intensity thresholds (**Fig. 4**) and the poor decoding accuracy observed at higher tone frequencies (**Fig. 7A**) under anesthesia. More complex spectral patterns (chords, filtered noises, complex sounds and decomposed complex sounds) were also better decoded with awake data (**Fig. 7C**). Slow temporal features of sound intensity ramps and frequency chirps were decoded in both states with similar efficiency. However, the fast sinusoidal variations of amplitude modulated sounds (AM) were better decoded under anesthesia (**Fig. 7C**). This suggests that the neural code in the cochlear nucleus under anesthesia loses in dynamic range but also gains in temporal precision at fast time scales. Therefore, isoflurane anesthesia produces two main effects in the cochlear nucleus: (i) a redistribution of sound responsive neurons which we expect to strongly impair normal downstream decoding (**Figs. 5, 6**). (ii) a slight modification of sound information carried by the network, with sharpened tuning curves but poorer encoding of low intensity and high frequency sounds while fast temporal precision is slightly improved (**Figs. 7**).

**Figure 7:**
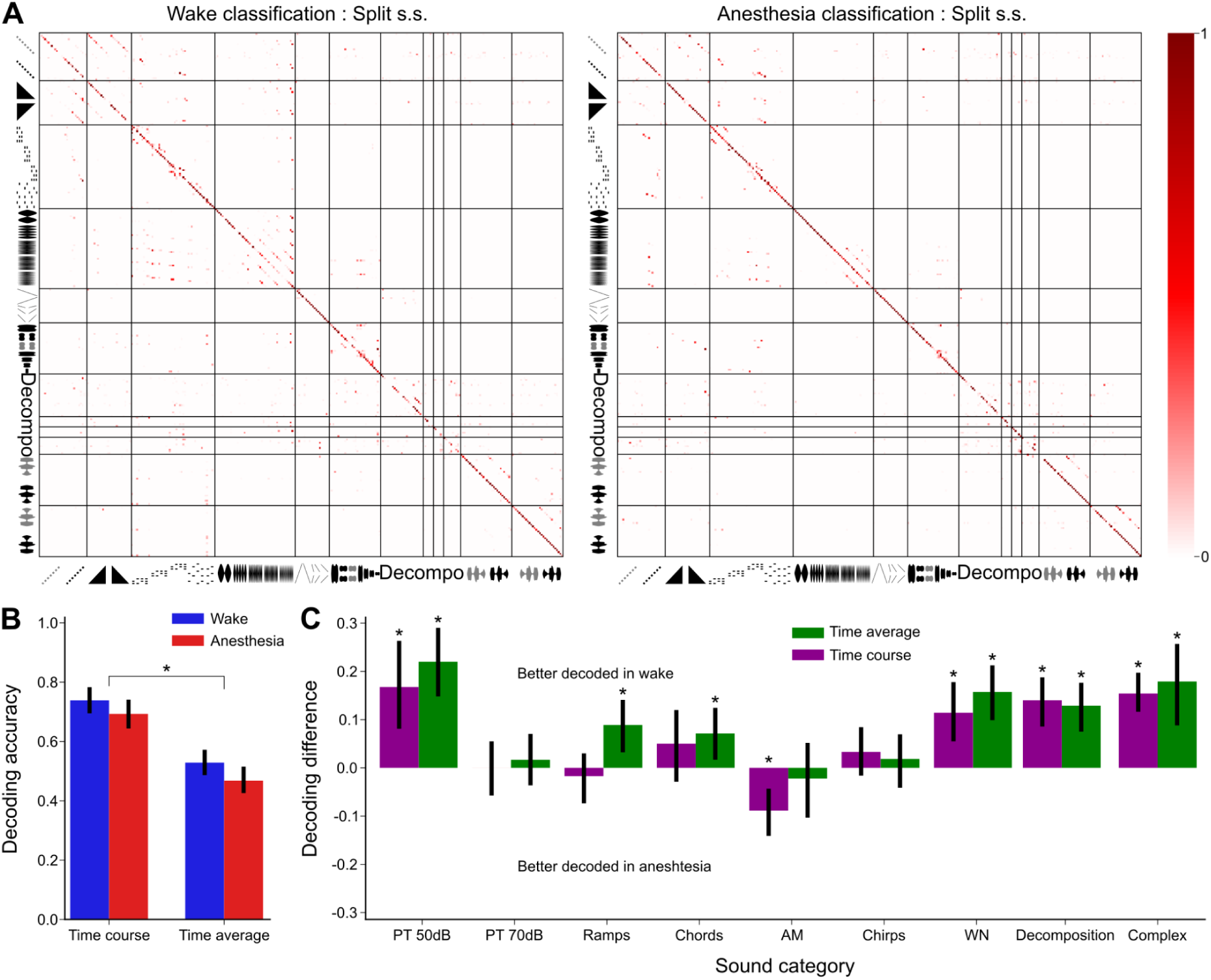
Isoflurane anesthesia differentially impacts the codes for distinct acoustic cues. **A.** Matrices of nearest-neighbors decoding algorithm output based on responses of neurons to sounds without time average, during the awake (left) and anesthetized (right) procedures. The color of the squares indicates the probability of the decoder to classify the input sound (in rows) into an output sound (in columns). A redder diagonal thus indicates a better decoding, whereas off-diagonal red pixels indicate the classifier confuses the two sounds. **B.** Barplots of same-state classification accuracy for each category in split s.s. dataset with the full time course of the responses (left) or after time-averaging the neural response (right). In both states, classification is significantly better using the full time course of neuron responses (Wake time course accuracy=0.72±0.02, time average accuracy=0.53±0.02, p=3e-13. Anesthesia time course accuracy=0.69±0.02, time average accuracy=0.49±0.02, p=1e-15, Mann-Whitney U test). **C.** Barplots of the difference of decoding accuracy in the split s.s. dataset between wake and anesthesia, for both time course and time average classification. Error bars correspond to the 5th and 95th percentiles of the bootstrap distribution (100 resampling). P-value is computed as the percentage of values above 0 -or below for categories with negative value of the mean- (PT 50dB time average: p<0.01, time course: p<0.01. PT 70dB time average: p=0.29, time course: p=0.47. Ramps time average: p<0.01, time course: p=0.31. Chords time average: p=0.01, time course: p=0.12. AM time average: p=0.37, time course: p<0.01. Chirps time average: p=0.23, time course: p=0.11. WN time average: p<0.01, time course: p<0.01. Decomposition time average: p<0.01, time course: p<0.01. Complex time average: p<0.01, time course: p<0.01).

## Discussion

Following for the first time the same neurons across wakefulness and isoflurane anesthesia in a peripheral sensory structure, the cochlear nucleus, which processes all auditory information arriving in the brain, we observed that anesthesia induces major changes in early auditory processing. The first modification is the >3-fold reduction of spontaneous activity levels under anesthesia (**Fig. 1**). Although the evaluation of population firing rate modulation through the magnitude of high frequency modulation gives only a rough estimate, the large effect clearly indicates that anesthesia brings the cochlear nucleus network in a different dynamic state compared to wakefulness. Isoflurane is known to boost inhibitory currents by increasing the efficiency of GABA_A_ receptors (*41*, *42*). The strong reduction in firing rate may therefore reflect reinforced local inhibition. The cochlear nucleus is a complex circuit with specificities in each of its subdivisions (*43*–*45*). However, most subdivisions have feedforward (*45*–*47*) and/or feedback inhibition (*48*). The increased potency of inhibitory synapses could lead to an increased baseline inhibition and thereby to the observed reduction of spontaneous activity (**Figs. 1 & 3**) (*49*). Given that the auditory nerve is constantly active (*50*), feedforward inhibition could have a non-negligible role, although it is unknown if isoflurane also impacts the inner ear and its baseline activity. Increased baseline inhibition could also be the mechanism for the elevation of sound intensity threshold observed both in single neurons (**Fig. 4**) and in auditory brainstem responses (ABR) (*51*) under anesthesia. This would be compatible with the observation that anesthetics which produce less potentiation of inhibition, such as ketamine-xylazine, also have less effect on the ABR threshold (*52*). Nevertheless, given the complexity of the cochlear nucleus circuits these potential mechanisms have to be carefully evaluated.

In practice, the dramatic impact of isoflurane anesthesia on firing rates raises unique challenges for spike sorting algorithms which could not robustly isolate single units from both the awake and anesthetized states without state-specific adjustment of their parameters (**Fig. 2**, see also Methods). An impact of anesthesia on spike sorting has already been documented for other anesthetics in the auditory cortex (*37*). For the cochlear nucleus, we encountered a combination of issues related to the strong modification of neural population dynamics across states, with a much larger number of spikes in wakefulness (**Figs. 1 & 2**) and bidirectional modulations of firing rates across states in individual neurons (**Fig. 5**). The larger number of spikes in wakefulness disrupted spike sorting by biasing cluster discovery towards units active in wakefulness, leading to a large number of undetected units in anesthesia when performing spike sorting with the same parameters across both states. We corrected this issue by forcing spike clustering exclusively on anesthesia. We obtained units that were more active in wakefulness than under anesthesia and others that were more active under anesthesia. Lower activity under anesthesia cannot result from a sampling bias because spike detection is easier under anesthesia, due to the lower background noise (**Fig. 1F-H**). By contrast, units presenting higher activity under anesthesia may potentially result from more difficult detections of their spikes in wakefulness. We ruled out this possibility by performing high quality unit identification with the split spike sorting procedure (**Fig. 2**). This showed that strong upregulation of activity under anesthesia happened even in units whose waveforms clearly peak above the noise levels of both awake and anesthetized states (**Fig. 5, Supplementary Fig. 1**). Spike sortings may also have been impacted by potential changes in the spike waveform across states. Three factors may have changed the spike waveform. First, larger background noise can bias detections toward larger spikes, leading to subsampling of a subset of the waveforms in wakefulness. Second, larger firing rate impacts the intracellular and extracellular spike waveform of neurons potentially due to the differential recruitment of voltage-dependent channels (*34*). Third, it has also been proposed that the overall population firing level influences electrical conduction in the neuropil (*34*, *53*) which may lead to changes in spike waveforms across states. Together, these effects likely explain the differences in the spike waveforms observed between anesthesia and wakefulness in some of the tracked single units (**Fig. 2E**, **Supplementary Figs. 1 and 2**). However, our analysis of spike waveform similarity between states shows that the changes observed in spike waveforms are weaker than the variability expected from chance similarity (**Fig. 2E**, **Supplementary Figs. 1 and 2**). Therefore, the modifications of sound responses under anesthesia do not result from matching errors.

The redistribution of sound-evoked activity we observed in the cochlear nucleus across wakefulness and anesthesia has two fundamental implications. First, it indicates that the representations produced by an early brainstem relay of auditory information result not only from feedforward propagation of upstream signals but also from rich network interactions. The balance of these interactions can be sufficiently perturbed by an external modulation, such as isoflurane, to deeply remodel the representations. Therefore, the functional identity of a neuron identified in anesthesia does not necessarily reflect its identity in wakefulness (**Fig. 5**). A similar shift of functional identities was recently observed under anesthesia in the auditory cortex (*6*). However, in the cortex, this shift is combined with a massive drop of the sound information carried by the population, and a convergence of spontaneous and evoked activity patterns. Neither of these effects were observed in the cochlear nucleus. The level of sound information carried by cochlear nucleus populations of similar sizes is similar across states (**Fig. 5**). Moreover, spontaneous activity in the cochlear nucleus is drastically reduced and does not resemble evoked activity (**Fig. 3**). This is in line with the view that the convergence of spontaneous and evoked activity under anesthesia is a cortex-specific process, not observed in the auditory thalamus (*6*) or brainstem (**Fig. 3**). The second important implication of our results is that the remodeling of auditory representations by anesthesia in the early auditory system likely impairs normal downstream integration of sensory information. This is clearly indicated by the very poor cross-state decoding performance observed for populations of neurons tracked across states (**Fig. 6**). Neurons of the downstream targets of the cochlear nucleus build their receptive fields based on precise combinations of synaptic inputs. If these inputs drastically change the magnitude and sign of their response in an heterogeneous manner (**Fig. 5**), the selectivity of the recipient neuron is bound to be modified. Although we cannot fully rule out the existence of complex compensatory mechanisms or the possibility that feedforward propagations entirely relies on the small fraction of neurons unaffected by anesthesia in the cochlear nucleus, the expected consequence of the large impact of anesthesia in the cochlear nucleus is a deterioration of downstream representation in the inferior colliculus, thalamus and cortex. Such a deterioration is indeed seen in the last two structures (*6*). Because the modifications of response properties in the cochlear nucleus affect the sign and magnitude of responses rather than the frequency selectivity, our results are compatible with the preservation of tonotopy at the mesoscopic scale across the auditory system under anesthesia (*7*, *54*, *55*). Yet we expect a strong impact on the microscopic organization and on the tuning to more complex acoustic patterns than simple pure tones. Given the brain-wide impact of anesthetics, many other processes likely contribute to the deterioration of auditory representation in the late auditory system. However, our observation challenges the view that anesthesia mainly impacts processing in higher order processing centers. On the contrary our results demonstrate that peripheral sensory networks, which are already complex circuits, are sufficiently perturbed to compromise normal information transfer upstream.

## Material and Methods

### Subjects and ethical authorizations

All mice used for electrophysiology were 10 to 12 weeks old male C57Bl6J mice (26-27g) that had not undergone any other procedures. Mice were group-housed (2–4 per cage) before and after surgery, had ad libitum access to food and water and enrichment (running wheel, cotton bedding and wooden logs) and were maintained on a 12-hour light-dark cycle in controlled humidity and temperature conditions (21-23°C, 45-55% humidity). All experiments were performed during the light phase. All experimental and surgical procedures were carried out in accordance with the French Ethical Committee the French Ethical Committees #89 (authorizations APAFIS#27040-2020090316536717 v1).

### Surgical procedures and electrophysiological recordings

To access the cochlear nucleus in awake mice, we performed an initial surgery during which we exposed the bone above the dorsal cerebellum and positioned a head-post for reliable stereotaxic placement of the mouse head in the electrophysiology recording apparatus. Mice were injected with buprenorphine (Vétergesic, 0,05-0,1 mg/kg) 45 min prior to surgery. Induction of anesthesia was carried out using 3% isoflurane. After induction, mice were kept on a thermal blanket with their eyes protected with Ocrygel (TVM Lab), and anesthesia was maintained with 1.5% isoflurane delivered via a mask. Lidocaine was injected under the skin 5 minutes prior to incision. The skull above the inferior colliculus and cerebellum was exposed for ulterior craniotomy. A well was formed around it using dental cement in order to retain saline solution during recordings and the head post was fixed to the skull using cyanolit glue and dental cement (Ortho-Jet, Lang). To protect the skull, the well was filled with a waterproof silicone elastomer (Kwikcast, WPI) that could be removed prior to recording. After surgery, mice received a subcutaneous injection of 30% glucose and metacam (1 mg/kg) and subsequently housed for one week with metacam delivered via drinking water or dietgel (ClearH20).

After recovery, mice were trained to remain quietly head-fixed for four days before recording by keeping them head-restraint for 30 min on day 1 up to 2 hours on day 4. The day before recording, mice were briefly anesthetized using isoflurane 1.5% in order to perform craniotomy and durectomy for electrode insertion. Following this training, a craniotomy and a durectomy were performed above the cerebellum and inferior colliculus in a brief surgery under isoflurane anesthesia. After at least one night of recovery, the awake mouse was head-fixed and Neuropixels 1.0 probes (384 channels) were inserted at 38-40° angle through the cerebellum, targeting the contralateral cochlear nucleus. Electrode angle and entry point were defined relative to the initial head-post placement (**Fig. 1A, B**). Fine tuning of these targeting parameters was progressively obtained through repeated penetrations based on time-locked responses to sounds easily detectable during probe insertion. For post-hoc histological verification of the electrode track using fluorescent dye, the electrodes were dipped in diI, diO or diD (Vybrant™ Multicolor Cell-Labelling Kit, Thermofisher) prior to insertion. Recordings were performed using warmed saline filling the cyanolit glue well and in contact with the reference electrode. After each recording the well was amply flushed and then refilled with Kwickast. Data was sampled at 30kHz using a NI-PXI chassis (National Instruments) and the SpikeGLX acquisition software. Recording could be repeated up to 3 days in a row to perform stable extracellular recordings. Using this approach, we recorded extracellular neuronal activity in the cochlear nucleus during 7 recording sessions in 4 mice. 5 recordings targeted the postero-ventral cochlear nucleus (PVCN) and 2 the dorsal cochlear nucleus (DCN), as defined based on post-hoc histology (**Fig. 1B**).

### Sound set and experimental protocol

Sounds were generated with Python (The Python Software Foundation, Wilmington, DE) and were delivered at 192 kHz with Matlab (The Mathworks, Natick, MA), using a NI-PCI-6221 card (National Instruments) driven by a custom protocol using the Matlab Data Acquisition toolbox and feeding an amplified free-field loudspeaker (SA1 and MF1-S, Tucker-Davis Technologies, Alachua, FL) positioned in front of the mouse, 10 to 15 cm from the mouse ear. Sound intensity was cosine-ramped over 10 ms at the onset and offset to avoid spectral splatter. The head fixed mouse was isolated from external noise sources by sound-proof boxes (custom-made by Decibel France, Miribel, France) providing 30 dB attenuation above 1 kHz.

We subsequently presented two sets of sounds to the animal. The first set consisted of 307 short sounds (<500ms, sketched in **Fig. 1A**) each repeated 12 times and played in a random order with a 1 s interval between sound onsets in 123 blocks of 30 sounds. 28 Pure tones: pure tones at 14 frequencies logarithmically spaced between 2 kHz and 60 kHz at 50 dB and 70 dB. 26 Ramps: up- and down-intensity ramped sounds at the same frequencies as the 13 pure tones between 2 kHz and 50 kHz between 50 dB and 70 dB. 48 Chords: summation of 2-4 70dB pure tones from low, medium, high frequency groups (11×3 sounds), broadly sampled frequencies (10 sounds) and harmonically arranged frequencies (5 sounds). 20 Chirps: up- and down- frequency sweeps of different durations between 25 ms and 400 ms at 6-12 kHz, 50 dB (10 sounds) or different frequency content between 4 kHz and 50 kHz at 50 dB in 500 ms (10 sounds). 30 Coloured noises (WN): broadband noises at 50 dB, 70 dB or up-/down-ramped in 100 ms and 500 ms (6 sounds), filtered noises at different bandwidths between 2 kHz and 80 kHz (14 sounds), and summation of two 1 kHz bandwidth filtered noises up- and down-ramped between 50 dB and 70 dB, matching frequencies of a subset of chords (10 sounds). 48 amplitude-modulated sounds (AM): AMs at 6 different modulation frequencies between 4 Hz and 160 Hz and 8 carrier frequency contents (2 pure frequencies, 5 sums of frequencies matching chords and 1 broadband noise). 60 Complex sounds: 15 complex sounds (recordings of accelerated music, animal calls, and natural environments) high-pass filtered at 2 kHz, played forward or time reversed, at 50 dB and 70 dB. 47 Decomposition sounds: snippets extracted from 4 selected complex sounds (2 bird calls, 1 dolphin call, 1 natural environment), and their composition which reconstruct their associated complex sound. The second set consisted of ten samples of natural sounds (30s long - two recordings in a cage containing a mouse family ∼10 days after birth, four recordings of urban cafés and street sounds, and four recordings from forest, ice, and country sounds) played twice each, interleaved between random blocks of the short sounds, which was not used in this study. These soundsets were presented first to the awake mouse (80 minutes), then rapid anesthesia was induced by applying a nose mask with a flow of isoflurane 2%. The isoflurane concentration was then decreased to 1.1% corresponding to a light narcosis state, ∼0.1-0.2% above the point at which spontaneous whisking could be observed. The continuity of anesthesia over the full procedure was monitored using infrared video of the mouse head (**Fig. 1D**). A second presentation of the soundset was performed under anesthesia, then the mask was removed for the mouse to wake up.

### Histology

In order to extract the brain for histology, mice were deeply anesthetized using a ketamine-medetomidine mixture and perfused intracardially with 4% buffered paraformaldehyde fixative. The brains were dissected and left in paraformaldehyde overnight and then sliced into hundred micrometer sections using a vibratome and mounted. Analysis of the fluorescence band diI, diO or diD allowed isolating up to 3 tracks per mouse for electrophysiological experiments.

### Data preprocessing and spike sorting

Raw data were band-pass filtered (300 - 6 000 Hz) and channels from the electrode tip (corresponding to the cochlear nucleus region) were selected using SpikeInterface (https://github.com/SpikeInterface). Isolated clusters were identified using Kilosort 2.5 followed by manual curation based on the interspike-interval histogram and the inspection of the spike waveform using Phy (https://github.com/cortex-lab/phy). Canonical spike sorting was first applied with common parameters throughout the whole recording, attempting to optimize the spike detection and assignment to clusters. As documented in the results and **Fig. 2**, no set of parameters could cluster spikes detected in anesthesia because of the drastic change in spiking activity it induced. We therefore designed two alternative procedures: a first aimed at maximizing the quality of single units clustered (split spike sorting) and a second aimed at maximizing the continuity of clusters between wake and anesthesia (consensus spike sorting). For split spike sorting, two separate spike sortings were conducted on the awake and anesthetized procedures, each with an optimized set of parameters. Due to the low spiking activity during the anesthetized procedure, spikes could be more easily matched, therefore we could apply more stringent spike sorting parameters (detection threshold = 6, clustering threshold = 8, and matching thresholds = [11,8]. On the contrary, the background noise due to high spiking activity in wake demanded looser parameters for efficient sorting (detection threshold = 6, clustering threshold = 6, and matching thresholds = [10,6]). For the consensus spike sorting, some steps of the standard kilosort 2.5 algorithm were modified to use different parameters on the wake and anesthesia batches of the recording. Spike detection threshold was lower for anesthesia batches to account for higher noise levels computed during wake (4 for anesthesia versus 6 for wake), templates were learned only on spikes detected during anesthesia batches to avoid the overrepresentation of awake spikes (clustering threshold = 6), and template matching parameters were lowered for anesthesia spikes to match the lower detection threshold (matching threshold = [10,4] for wake, [10,4]*⅔ for anesthesia. After manual curation, single trial sound responses were extracted (0.3s before up to 1s after sound onset) as a histogram of 10ms time bin and the average activity over the prestimulus period (0.3s - 0s before sound onset) was subtracted for each trial. Based on histology, we identified a number of units whose location on the Neuropixel probe was not compatible with a localization in the cochlea nucleus.

### Matching of single units

Split spike sorting yielded two distinct sets of single units for the awake and anesthetized procedures of each experiment. Thus we identified post-hoc matching single units based on the correlation of their template. Each “awake” template was matched with the most correlated “anesthesia” template from the same experiment. We applied two criteria: the templates of the awake and anesthesia units must be more correlated than chance and they must be more strongly correlated with each other than any other unit. For the first criterion, a random distribution was constructed from the maximally correlated templates from different experiments. Based on this distribution, we inferred a correlation threshold (r=0.74) above which chance similarity had a probability lower than 0.05 (**Fig. 2E**). Selecting all units with waveform correlation > 0.74, we realized that several of them could be correlated with other units at a lower but yet similar level introducing an uncertainty in the matching of units. We therefore studied the distribution of waveform correlations between regularly spikesorted single units in all single session datasets and deduced our second criterion (no secondary match at a distance of <0.2 correlation), ensuring a low mismatch likelihood. Combining the two thresholds, we defined matched units as those having a waveform correlation ρ_*max*_ > 0. 74 across brain states and ρ < ρ_*max*_ − 0. 2 with any other simultaneously recorded unit. This stringent but necessary criteria allowed us to track 18 units across wakefulness. To visualize the similarity of spike waveforms with minimal impact of a potential drift of the probe, we compared the spike waveforms across the last 15 minutes of the awake procedure with the first 15 of the anesthetized one. For a few neurons with very low firing rates, we extended these time windows to include at least 100 spikes in both states. The majority of these waveforms were extremely similar across states (**Supplementary Fig. 1 & 2**).

### Neuron response properties

Spontaneous firing of neurons was computed during the 142 inter-block intervals, each lasting at least 3s. A window of 2s starting 0.3s after the offset of the last sound of a block was defined, during which all detected spikes were considered due to spontaneous activity of the neurons. The spontaneous firing rate in spikes per second was then defined as the sum of all detected spikes divided by the number of inter-blocks intervals multiplied by their duration (142*2). The mean activity evoked by sound presentation was defined as the average firing rate during sound presentation of all sounds minus the average firing rate during the 300ms baseline period before sound presentation. To estimate the broadness of frequency tuning, we computed the half-width response on the responses of neurons to 50dB and 70dB pure tones. This corresponds to the width of frequency around the peak of maximum response at which the neuron response is superior or equal to half its maximum (in octaves):

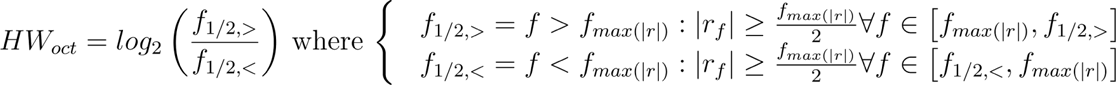

The precise location of half-peaks was estimated using a linear fit between sampled frequency responses. To estimate the intensity threshold of response of neurons, we used the response of neurons to sound ramping in intensity between 50 and 70dB at their preferred frequency *f*_*max*(|*r*|)_. The intensity threshold corresponds to the intensity of the sound at the time when the neuron firing rate exceeds twice the standard deviation of its spontaneous firing rate:

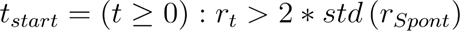

### Population activity classifiers

To evaluate the accuracy of sound identification based on single-trial population responses, we trained a nearest-neighbor classifier to categorize sounds based on the population response of neurons. For each state, population responses were estimated after pooling all neurons from all recording sessions into a pseudo-population. For train and test sets, a population vector was constructed from neuronal responses in each state for every sound, averaging over their respective subset of trials. All classifications were conducted on the full time course of neuronal response, except when stated otherwise (time average), in which case the neuronal response was average over the duration of sound presentation. 50 pairs of train-test population responses were created by randomly selecting half of the trials for each sound as a training set, and using the other half as the testing set. For each training set, the classifier takes as input a train sound population vector and outputs the test sound with the most correlated population vector. Accuracy of classification is defined as the average proportion of sounds correctly assigned by the classifier over all sounds. For ‘same state’ classification state, the training and testing sets were both defined on neural responses recorded during the same state, either in wakefulness or under anesthesia. For ‘cross state’ classification, the training set was defined on responses from one state, and the testing set on responses from the other state.

To assess the impact of anesthesia on the decoding of sound features, we computed the difference of decoding accuracy between wakefulness and anesthesia on different categories of sounds using the split s.s. dataset. To evaluate the statistical significance of the change in decoding, we used bootstrap procedure over neurons. 2×100 dataset of responses were created from random resampling of neurons in the awake and anesthetized datasets. The difference was computed over the classifier performance on all sounds of each category on those 100 bootstrap populations for 50 draws of train-test trial sets. The significance threshold was set at the 95th percentile of the bootstrap population.

### Principal component analysis and mouse state classification

Principal component analysis and classification of anesthetized vs awake states based on neural data were performed using the scikit-learn package for Python. The average response of neurons to sounds over time and trials were used as samples for computing the principal components. They were then projected into the first 3 principal components labeling each state with a different color. A linear singular vector classification was performed on the data projected into the component space that assigned the state from which sound responses were recorded. A 10-fold Cross-validation of the classifier was performed using a 1-to-1 ratio of sounds from each state in every fold.

### Statistical Analysis

The statistical tests used in this study are the Mann-Whitney U test for single distribution and unpaired distribution, the Wilcoxon signed-rank test for paired distributions, the Kruskal-Wallis for differences of distribution, and the bootstrap resampling test with 100 resamples for classifier comparisons. All tests are two-sided except the bootstrap test which is one-sided.

## Acknowledgments

We thank Maia Brunstein of the Hearing Institute Bioimaging Core Facility of C2RT/C2RA for help in acquiring histology images. We acknowledge the support of the Fondation pour l’Audition to the Institut de l’Audition.

## Funding

We acknowledge the support of the following funding sources: Fondation pour l’Audition, FPA IDA02 (BB) and APA 2016-03 (BB) European Research Council, ERC CoG 770841 DEEPEN, (BB)

## Author contributions

Conceptualization: EG, SB, BB

Methodology: EG, SB, BB

Investigation: EG

Data curation: EG

Formal analysis: EG

Supervision: BB

Project administration: BB

Funding acquisition: BB

Writing-original draft: EG, SB, BB

Writing-review & editing: EG, SB, BB

## Competing interests

Authors declare that they have no competing interests.

## Data availability

All datasets are freely available at DOI:10.5281/zenodo.10471543, hosted by Zenodo.

## Code availability

Custom codes used in this study are freely available at DOI:10.5281/zenodo.10471543, hosted by Zenodo.

## Notes

### Competing Interest Statement

The authors have declared no competing interest.

