## Supplementary Figures for "Massive perturbation of sound representations by anesthesia in the auditory brainstem"

Etienne Gosselin *et al.*

**This PDF file includes:**

Figs. S1 to S2

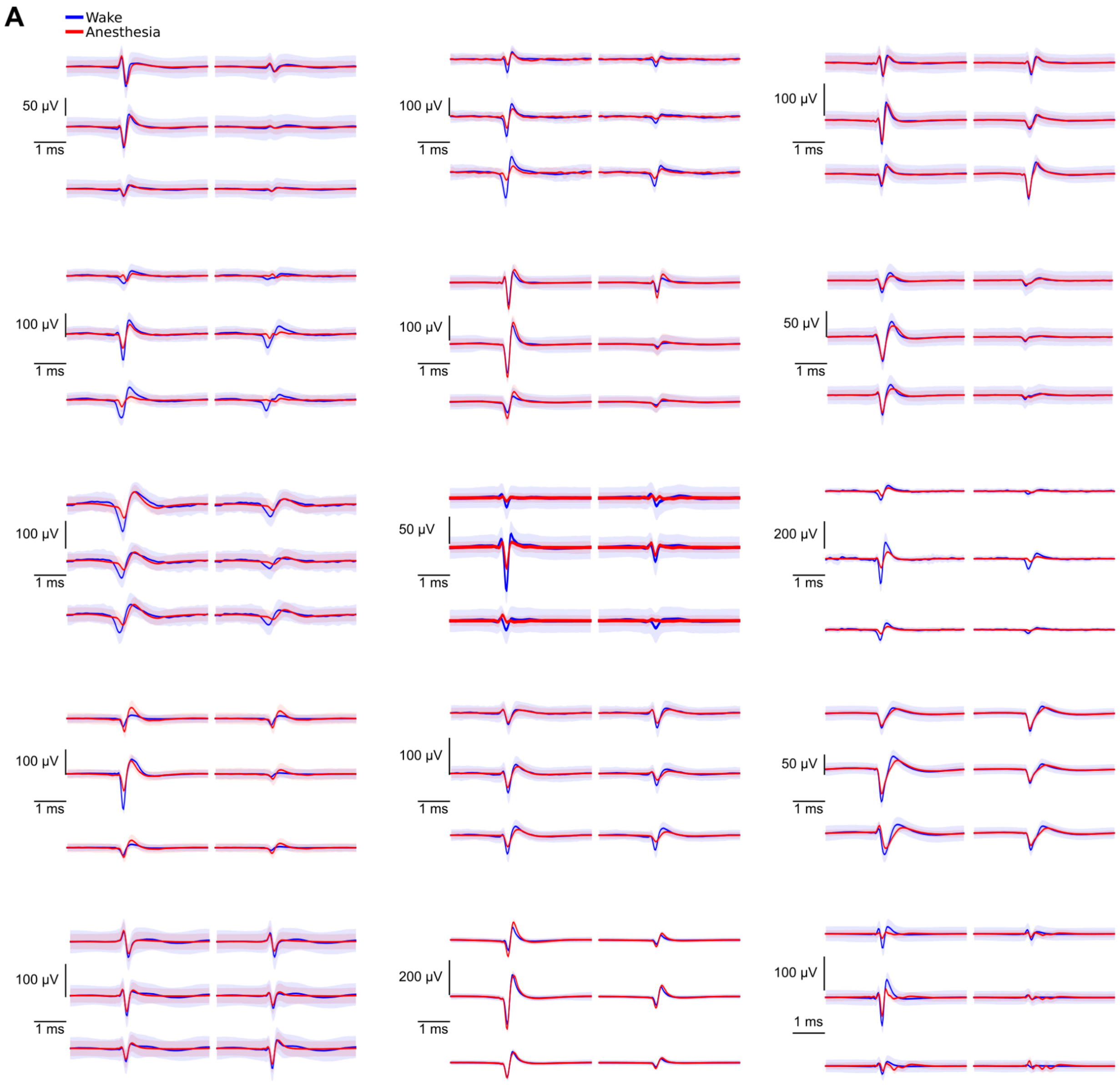


**Fig. S1.: Waveform similarity across anesthetized and awake states for matched units after split spike sorting**

**A.** Spike waveforms for 15 matched neurons of the split s.s. dataset for 6 channels centered around the maximal amplitude channel of each neuron.

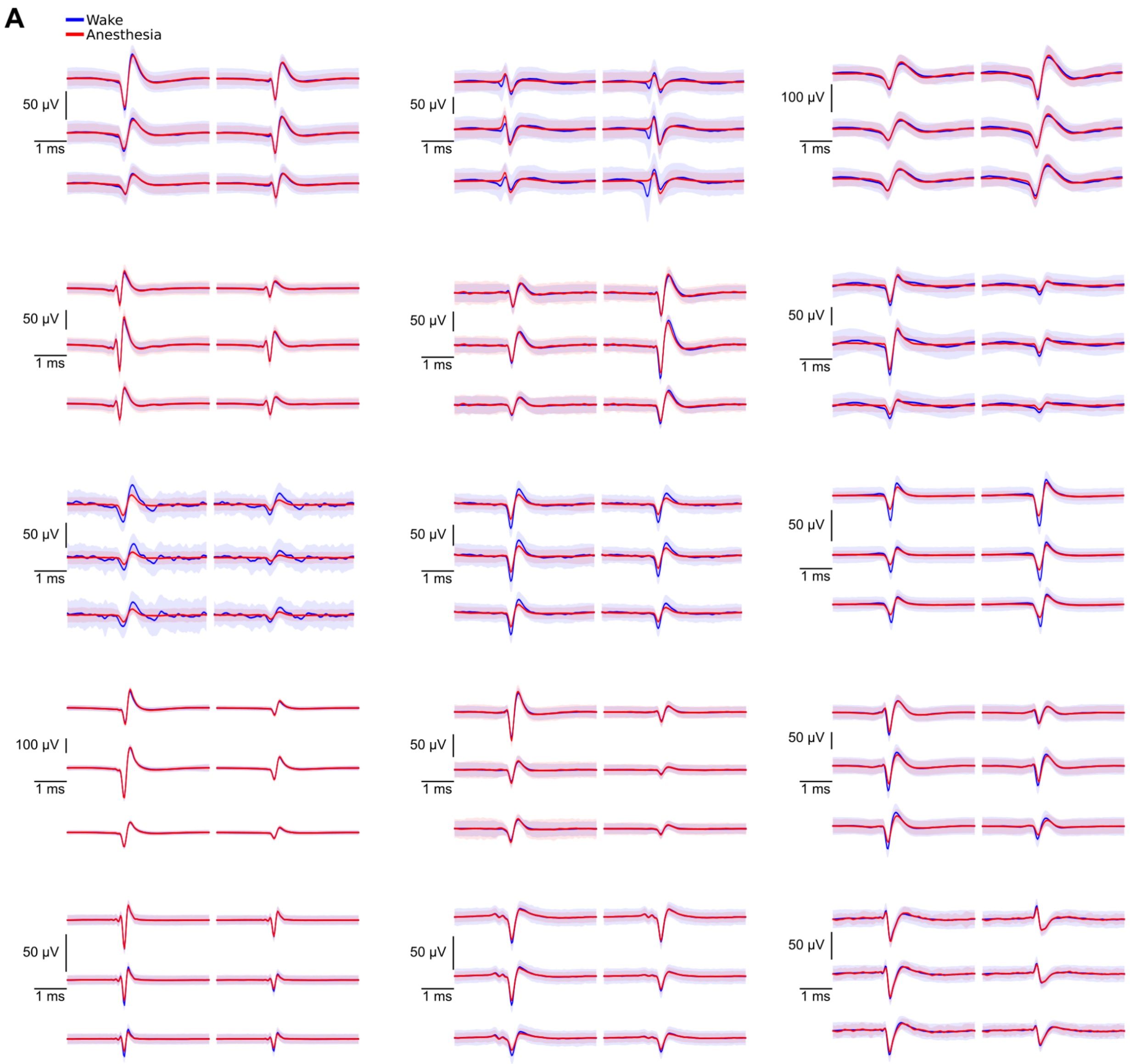


**Fig. S2.: Waveform similarity across anesthetized and awake states (consensus spike sorting)**

**A.** Spike waveforms for 15 matched neurons of the consensus spike sorting dataset for 6 channels centered around the maximal amplitude channel of each single unit.
